# Two new species of *Graphium* (Microascales, Ascomycota) associated with pine-infesting bark beetles in China

**DOI:** 10.1101/2024.09.14.613081

**Authors:** Jie Li, Xiang Gao, Kun Liu, Minjie Chen, Yutong Ran, Congwang Liu, Tong Lin, Mingliang Yin

## Abstract

*Graphium* is a genus of fungi that falls under the order Microascales of Ascomycota. Some species in this genus can establish a unique symbiotic relationship with the pine-infesting bark beetles, while others are typically found in wood or soil habitats. To comprehensively investigate the diversity of species of these fungi, recent field trips were conducted in seven provinces (Fujian, Guangdong, Guizhou, Guangxi, Liaoning, Shaanxi, and Shandong) in China. 96 pure isolates of *Graphium* were obtained by sequences from 361 samples. Nineteen representative strains were carefully selected to generate sequencing data from four gene regions (ITS, LSU, EF1A and TUBB), then used to construct phylogenetic trees for the genus. The results revealed the discovery of two new species, namely *G. armandii* sp. nov. and *G. massoniana* sp. nov., and *G. pseudoumiticum* was the most common species in various pine hosts.

## Introduction

The genus *Graphium* was established by Corda (1837), with *G. pennicillioides* as the type species (Hughes 1958). Because of its ecological niche and morphological similarity, this genus was once thought to be closely related to ophiostomatoid fungi (Upadhyay 1981, Seifert & Okada 1993). However, DNA sequence phylogenetic analyses accommodate *Graphium* within Microascales (Okada 1998). The sexual state of this genus is still unknown, whereas the asexual state usually produces broom-like, pigmented synnemata with annellidic sporulation and cylindrical hyaline conidia (Okada 2000). Some *Graphium* species often have a second anamorph, performing as scedosporium-like conidiophores (Jankowiak et al., 2023).

Currently, more than 140 legitimate species are registered on Mycobank, excluding subspecies or varieties, but only 20 species, most of which have been published with molecular data support. After the year 2000, all new species have DNA sequences, except for *G. variabile* and *G. wuweiense* (Wu et al. 2015). The commonly used barcoding gene segments within this genus are ITS and EF1A. Furthermore, the data volume of LSU and TUBB sequences is continuously increasing (Jankowiak et al. 2023).

Currently, three new species of the genus *Graphium* have been described in China, but none of them are related to bark beetles. One of them, *G. carbonarium*, was discovered in *Salix babylonica* infested by *Pissodes* sp. in Lijiang, Yunnan province (Paciura et al. 2010). The other two species were reported from different soil environments. *G. variabile* was isolated from forest soil in Xiamen, Fujian Province, and *G. wuweiense* was isolated from soil in a desert oasis in Wuwei, Gansu Province, respectively. (Wu et al. 2015).

Furthermore, three other known species (*G. fimbriisporum*, *G. laricis*, and *G. pseudormiticum*) were found associated with bark beetles in China. *G. fimbriisporum* was found to be associated with *Ips typographus* infesting *Pinus koraiensis* in Heilongjiang (Chang et al. 2019) and *G. laricis*, which has been found on larch trees (*Larix* spp.) damaged by *Ips subelongatus* in Inner Mongolia (Liu et al. 2016). Among these, *G. pseudormiticum* was considered the most commonly seen. This species was reported to be associated with *Pissodes* sp. in *Pinus yunnanense* (Cruywagen et al. 2010), *Ips acuminatus* infesting *P. keisya* in Puer, Yunnan Province (Chang et al. 2017), and *Cryphalus piceae* that infest *P. thunbergia* in Weihai, Shandong Province (Chang et al. 2021).

Between 2016 and 2022, several field trips were conducted in pine forests located in seven provinces of China. Fujian, Guangdong, Guizhou, Guangxi, Liaoning, Shaanxi, and Shandong. These field trips focused on studying bark beetle-associated fungi. Many graphium-like isolates were collected and preliminarily recognized during periods. This study aims to accurately identify these isolates by analyzing their morphological and molecular sequence data.

## Materials and Methods

### Fungal Isolation

The strains identified in this study were all from the Culture Collection of South China Agricultural University (SCAU). Ex-type cultures of new species were deposited in the Culture Collection of South China Agricultural University (SCAU) and the China General Microbiological Culture Collection Center (CGMCC). Type herbariums were preserved in the Fungarium (HAMS), institute of Microbiology, Chinese Academy of Sciences. Taxonomic novelties of undescribed species were registered in Mycobank (https://www. mycobank.org).

### DNA Extraction, PCR, and Sequencing

The isolates were cultured on MEA medium for 7-14 days to obtain mycelium for DNA extraction. About 100 mg of mycelium was transferred to a sterile 2 ml tube using an inoculation needle. DNA extraction was performed using the PrepMan® Ultra Sample Preparation Reagent (Applied Biosystems, USA).

Four gene regions (ITS, LSU, EF1A, and TUBB) were amplified for sequencing and phylogenetic analyses. The specific primers for the amplification of different gene segments are ITS5 and ITS4 for ITS (White et al. 1990), LR0R and LR5 for LSU, EF2F (Marincowitz et al. 2015), and EF2R (Jacobs et al. 2004) for EF1A, bt2a and bt2b for TUBB (Glass and Donaldson 1995).

The PCR reaction was carried out in a 25 μL mixture system, which included 1 μL of DNA template, 0.5 μL of each primer (10 μM), 12.5 μL of DreamTaq ^TM^ Green PCR Master Mix that contained Taq polymerase, MgCl_2_, dNTPs, and reaction buffer (Applied Biosystems in California, USA) and 10.5 μL of PCR grade water. Conditions for the PCR reactions were set as follows: an initial denaturation step at 95 ° C for 3min, followed by 35 cycles of 95 ° C for 30 s, 55-52 ° C for 30 s, and 72 ° C for 1 min, and a final elongation at 72 ° C for 10 min. The PCR products were detected by 1.5% agarose gel electrophoresis and sequenced by Sangon Biotech (Shanghai) Co., Ltd., Guangzhou Branch.

### Phylogenetic analyses

The consensus sequence of each strain was generated by assembling forward and reverse sequences in Geneious Prime v.11 software (Biomatters Ltd, Auckland, New Zealand). Subsequently, the consensus sequences were aligned with reference sequences using MAFFT 7 online (Katoh and Standley 2013). To ensure accuracy, alignments of all datasets were manually verified in MEGA 7 (Kumar et al. 2016), and sequences were submitted to GenBank (https://www.ncbi.nlm.nih.gov/genbank/).

All data sets (ITS, LSU, EF1A, TUBB) were subjected to maximum likelihood (ML), maximum parsimony (MP), and Bayesian inference (BI) analyses. The isolates of *Ambrosiella beaver* and *A. xylebori* represented the outgroup taxa.

ML estimations were conducted in MEGA X using the best substitution model, 1,000 bootstraps, and the nearest-neighbor-interchange (NNI) branch swapping option. The best substitution models were determined in JModeltest 2.1.10 (Posada 2008).

BI statistics were executed in MrBayes 3.2.6 (Ronquist et al. 2012) using Markov Chain Monte Carlo (MCMC) chains run from a random starting tree for five million generations. The trees were collected every 100 generations, and burn-in values were determined using Microsoft Excel 2021.

MP analyses were performed using PAUP* 4.0b10 (Swofford 2003), with 1000 replicates of heuristic searches and the tree bisection and reconnection (TBR) branch swapping options. Gaps were considered as the fifth base. Following tree generation, metrics such as tree length (TL), consistency index (CI), retention index (RI), homoplasy index (HI), and rescaled consistency index (RC) were documented.

### Morphology Characterization

Mycelium-inoculated agar blocks (3 mm × 3 mm) were placed next to sterilized pine twigs in the 2 % water agar medium (20 g agar in 1000 ml deionized water) and incubated at 25 ° C for 14 days. The reproductive structures of the isolates found in the twigs were transferred to water droplets on microscope slides using a syringe needle. The microstructures were then examined and captured under the microscope Axioscope 5, manufactured by ZEISS (Jena, Germany). Any new names of species were registered in Mycobank (www.mycobank.org).

## Results

### Phylogenetic analyses

A total of 96 isolates were obtained and preliminarily recognized as *Graphium* spp. Nineteen representative isolates used for phylogenetic analyses were listed in **Table 1**, and reference isolates of known species were listed in **Table 2**.

**Table 1.**
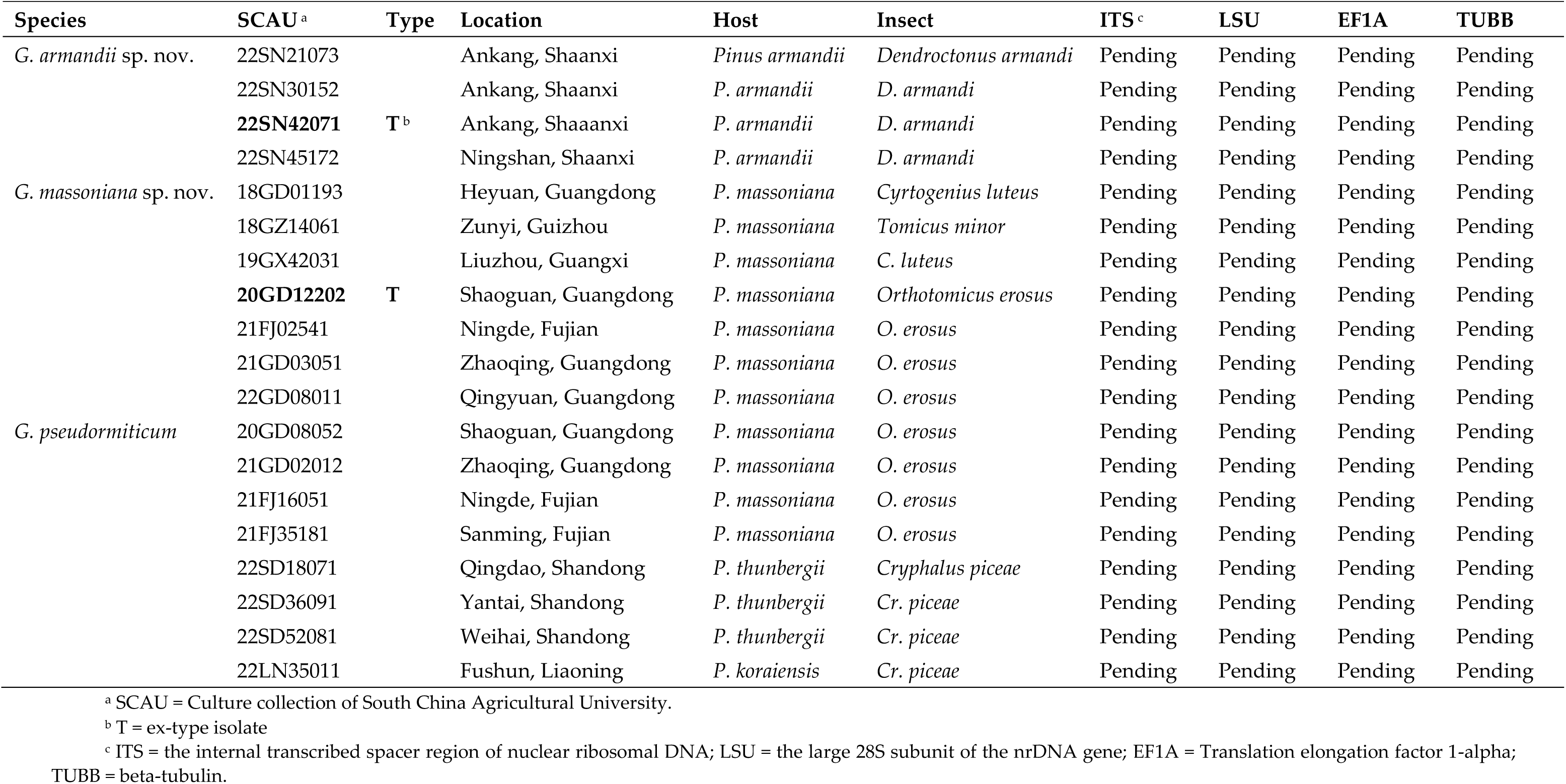
Representative isolates of *Graphium* collected in this study.

**Table 2.**
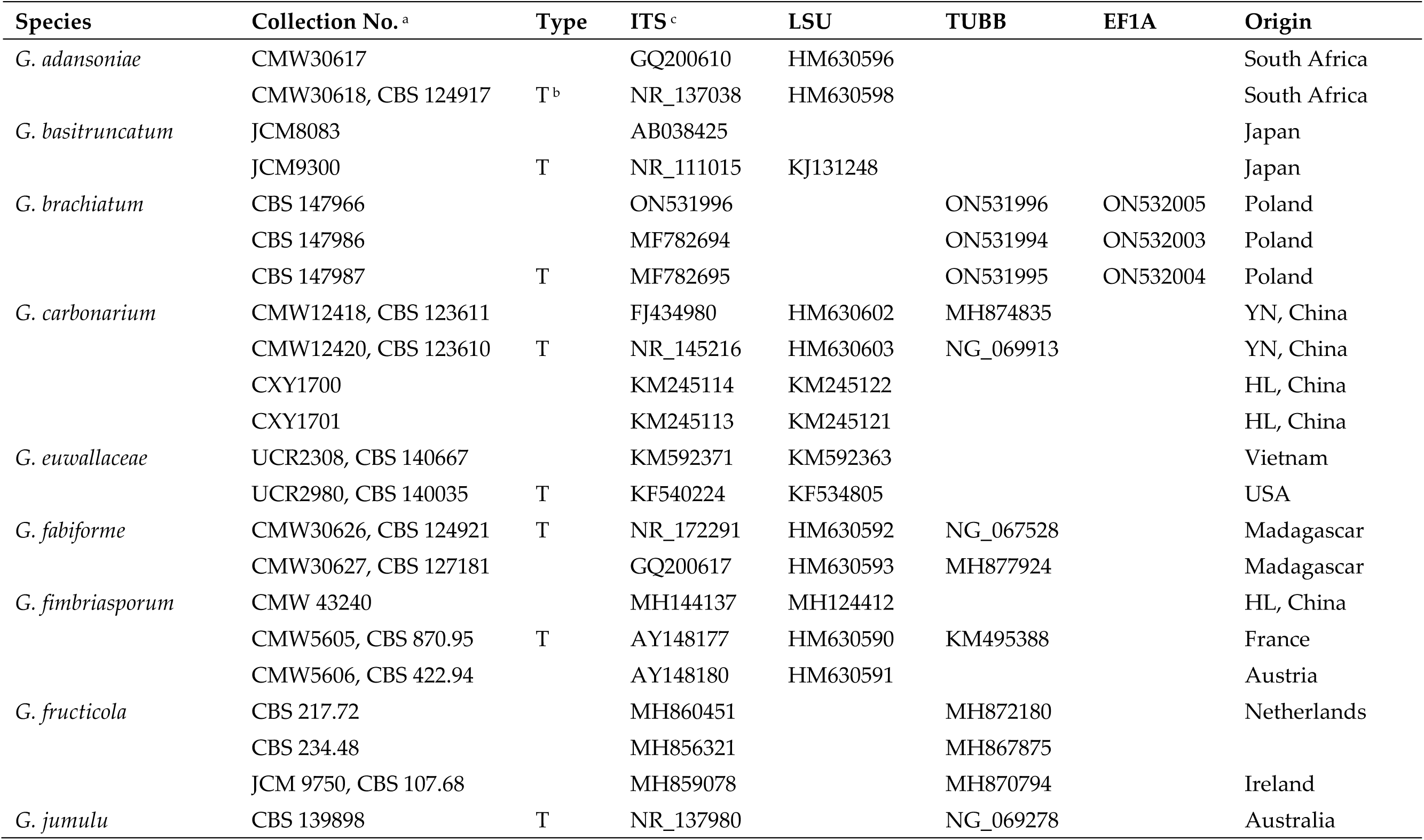
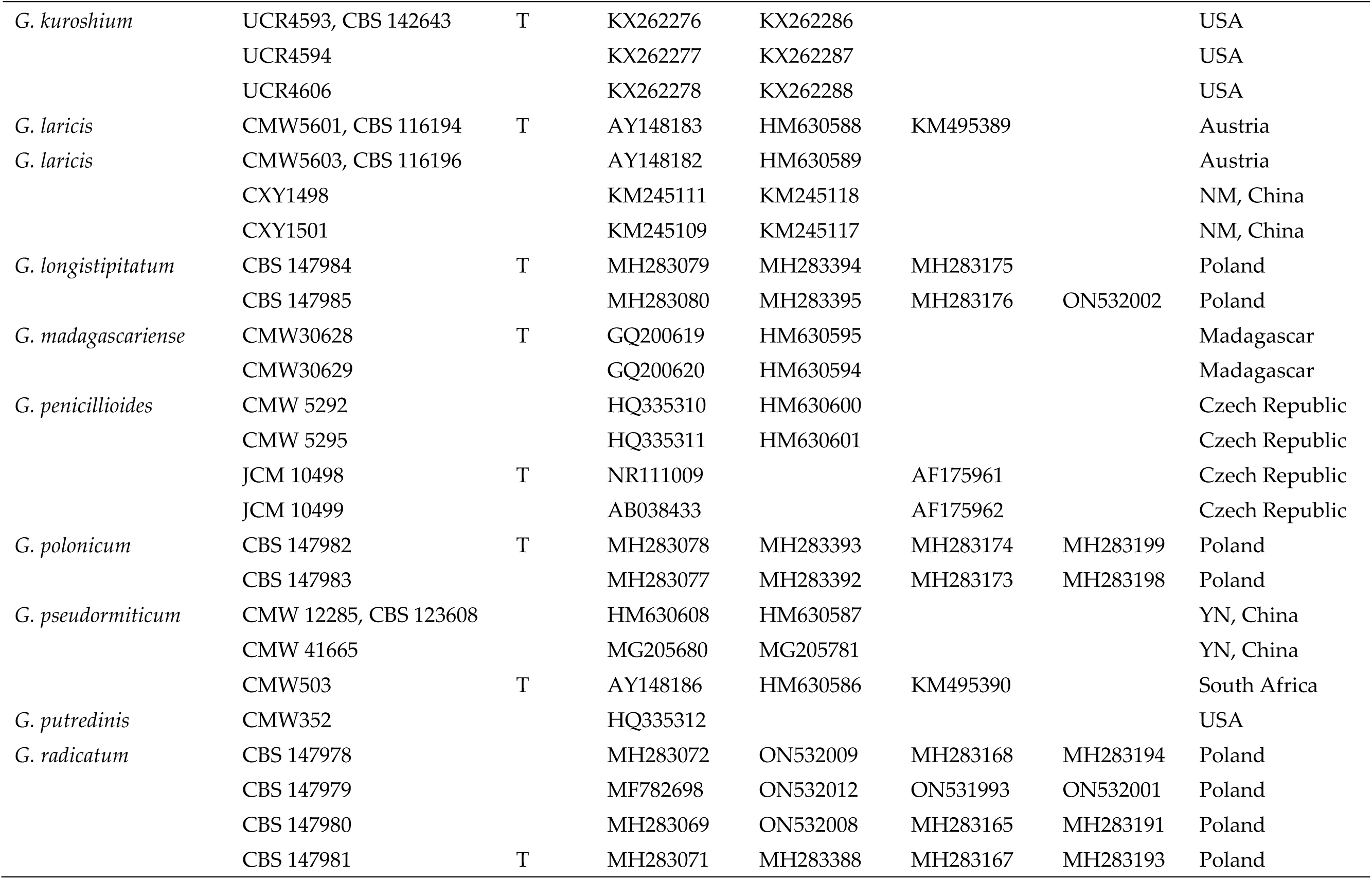
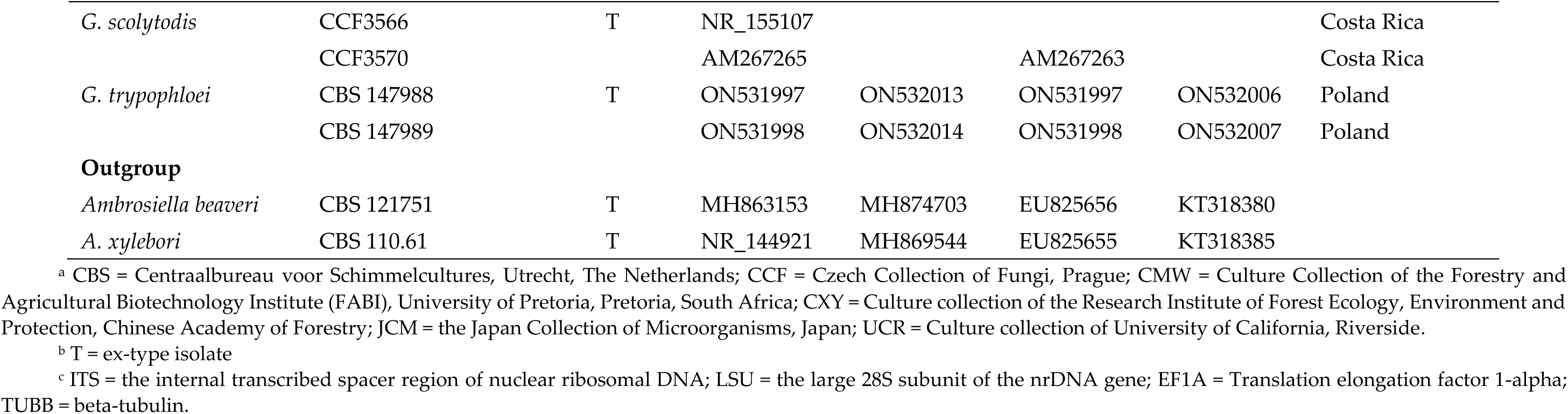
Reference isolates of *Graphium* used in this study.

The topologies of the trees obtained from the ML analyses were largely congruent with the bootstrap support values, as indicated in **Figs. 1, 2, 3, and** 4, respectively. The most important parameters used and the statistical values obtained from the phylogenetic analyses were listed in **Table 3**. The results showed that 19 representative isolates from this study formed three lineages in *Graphium*, including two undescribed species (Taxa 1 and 2) and one known species, *G. pesudomiticum* (Taxon 3).

**Figure 1.**
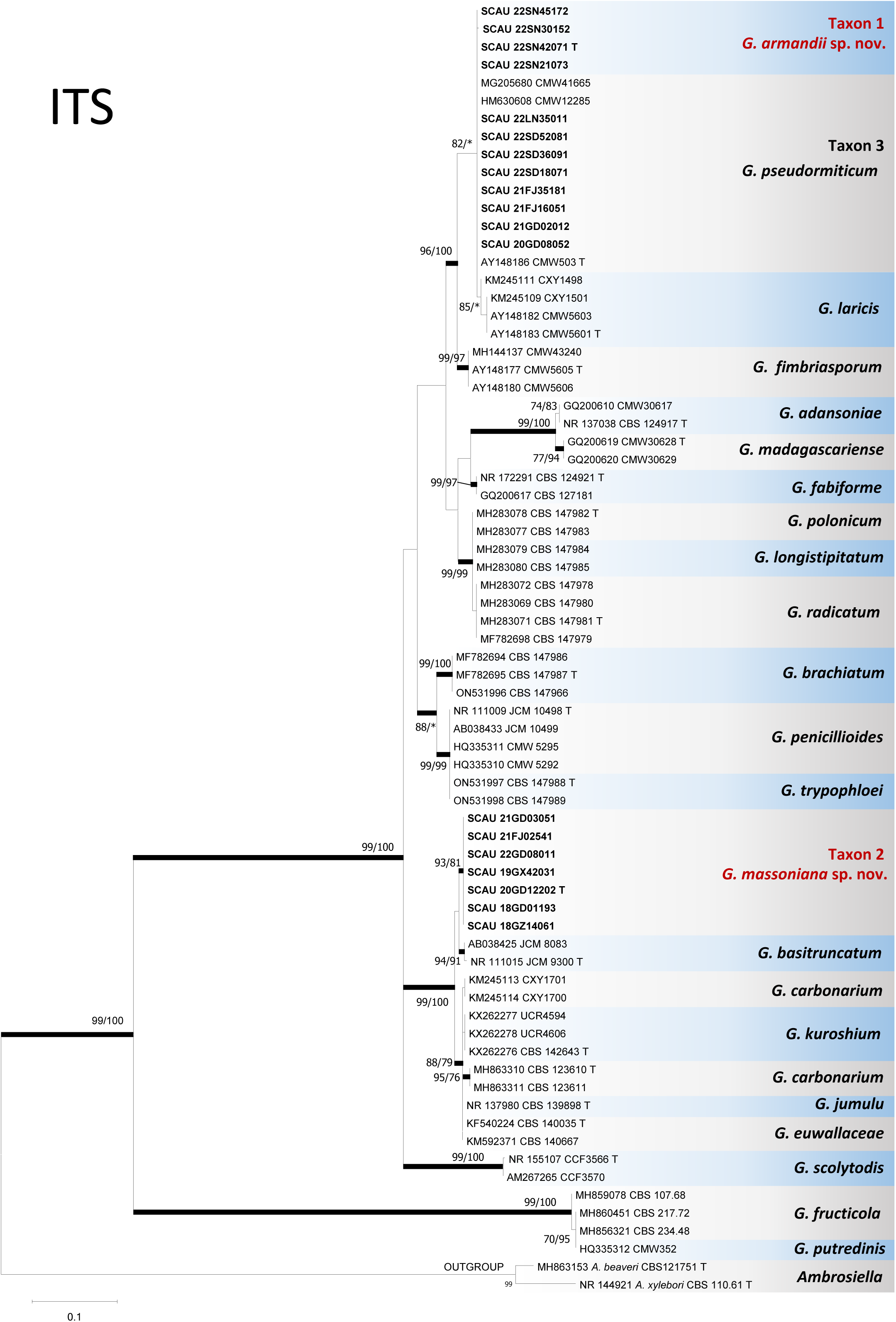
ML tree of *Graphium* generated from ITS sequence data. Sequences generated from this study are printed in bold type. Bold branches indicate posterior probability values ≥ 0.95. Bootstrap values ≥ 75 % are recorded at nodes as ML/MP. * Bootstrap values < 75 %. T = ex-type isolates.

**Figure 2.**
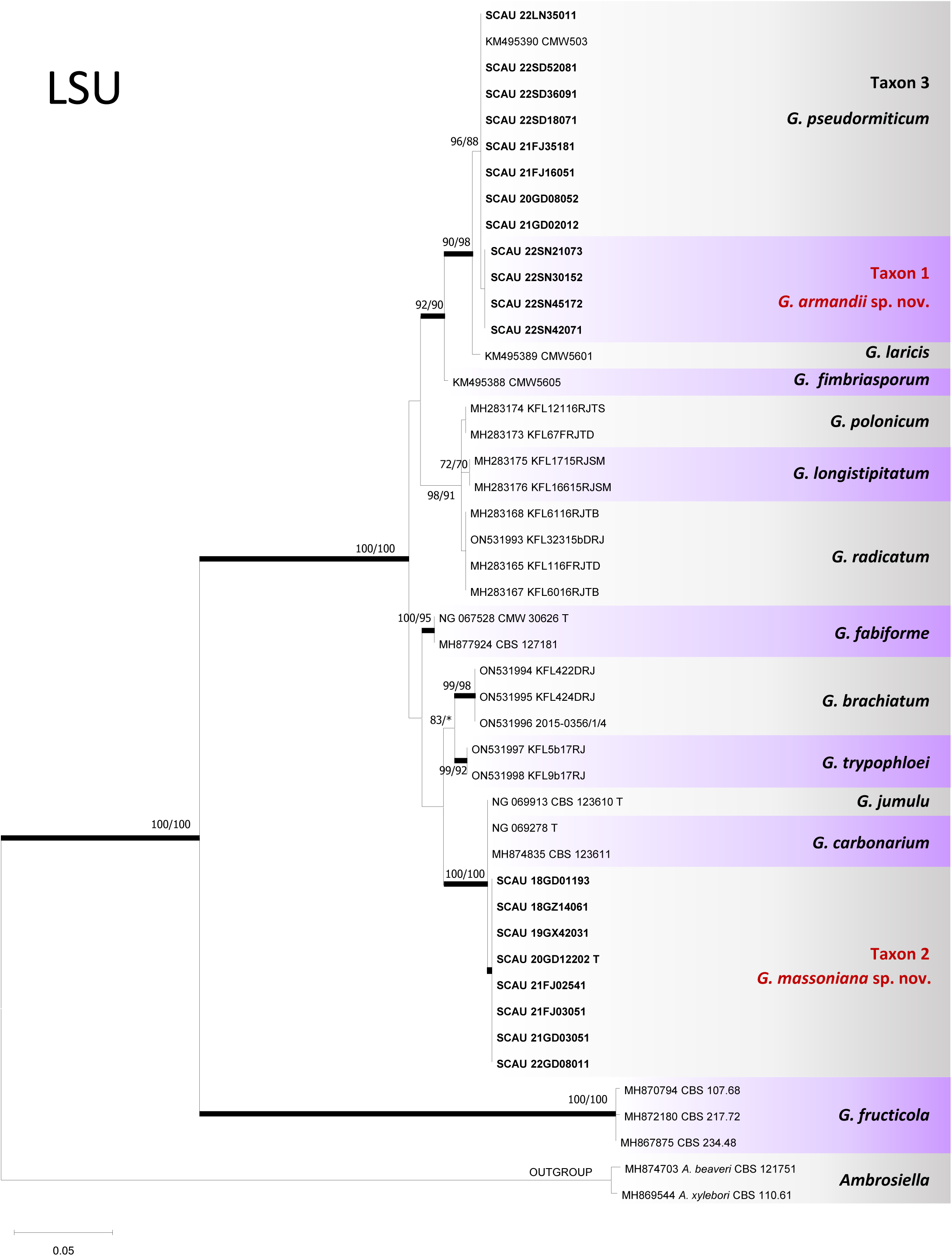
ML tree of *Graphium* generated from LSU sequence data. Sequences generated from this study are printed in bold type. Bold branches indicate posterior probability values ≥ 0.95. Bootstrap values ≥ 75 % are recorded at nodes as ML/MP. * Bootstrap values < 75 %. T = ex-type isolates.

**Figure 3.**
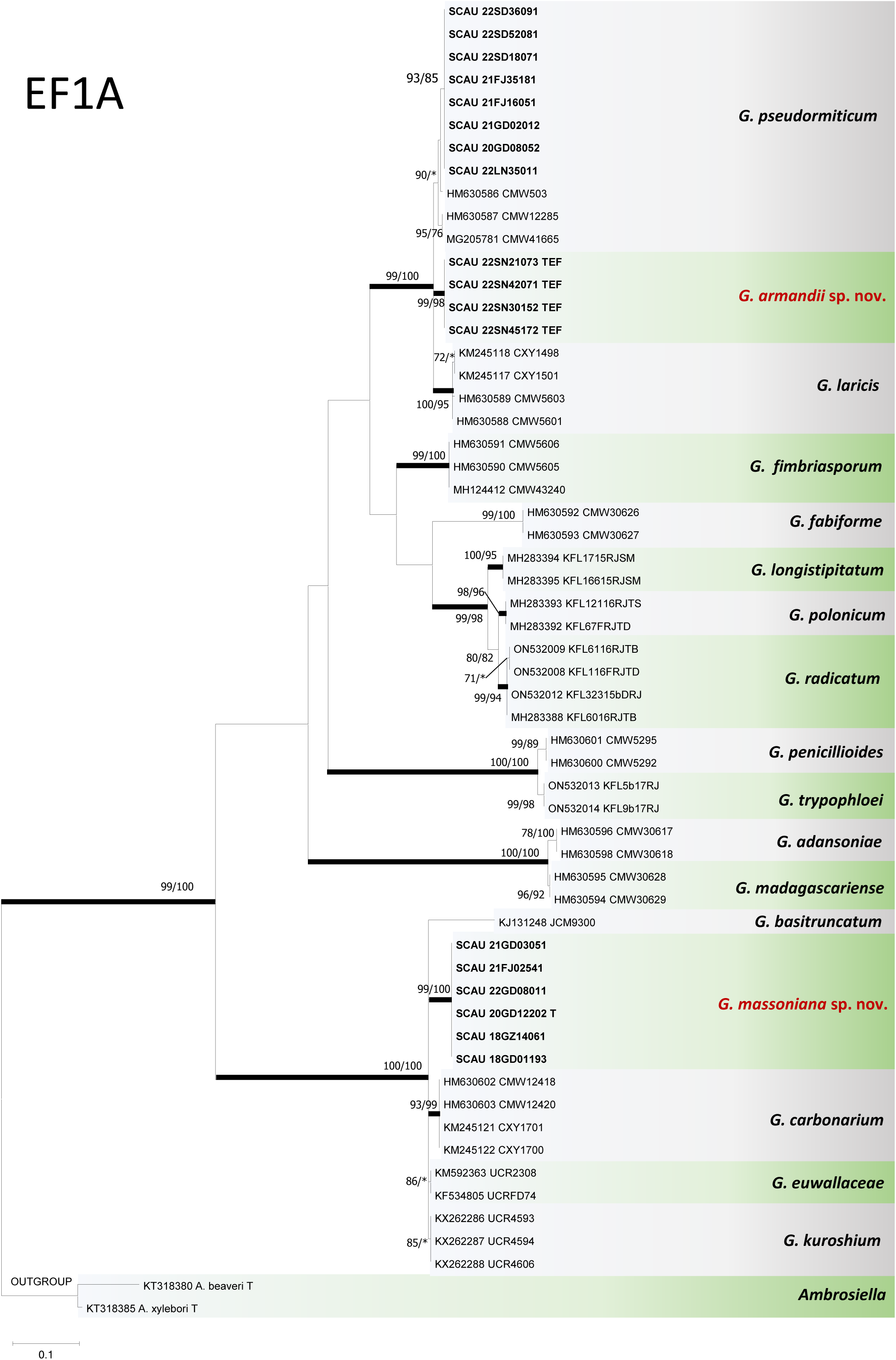
ML tree of *Graphium* generated from EF1A sequence data. Sequences generated from this study are printed in bold type. Bold branches indicate posterior probability values ≥ 0.95. Bootstrap values ≥75 % are recorded at nodes as ML/MP. * Bootstrap values < 75 %. T = ex-type isolates.

**Figure 4.**
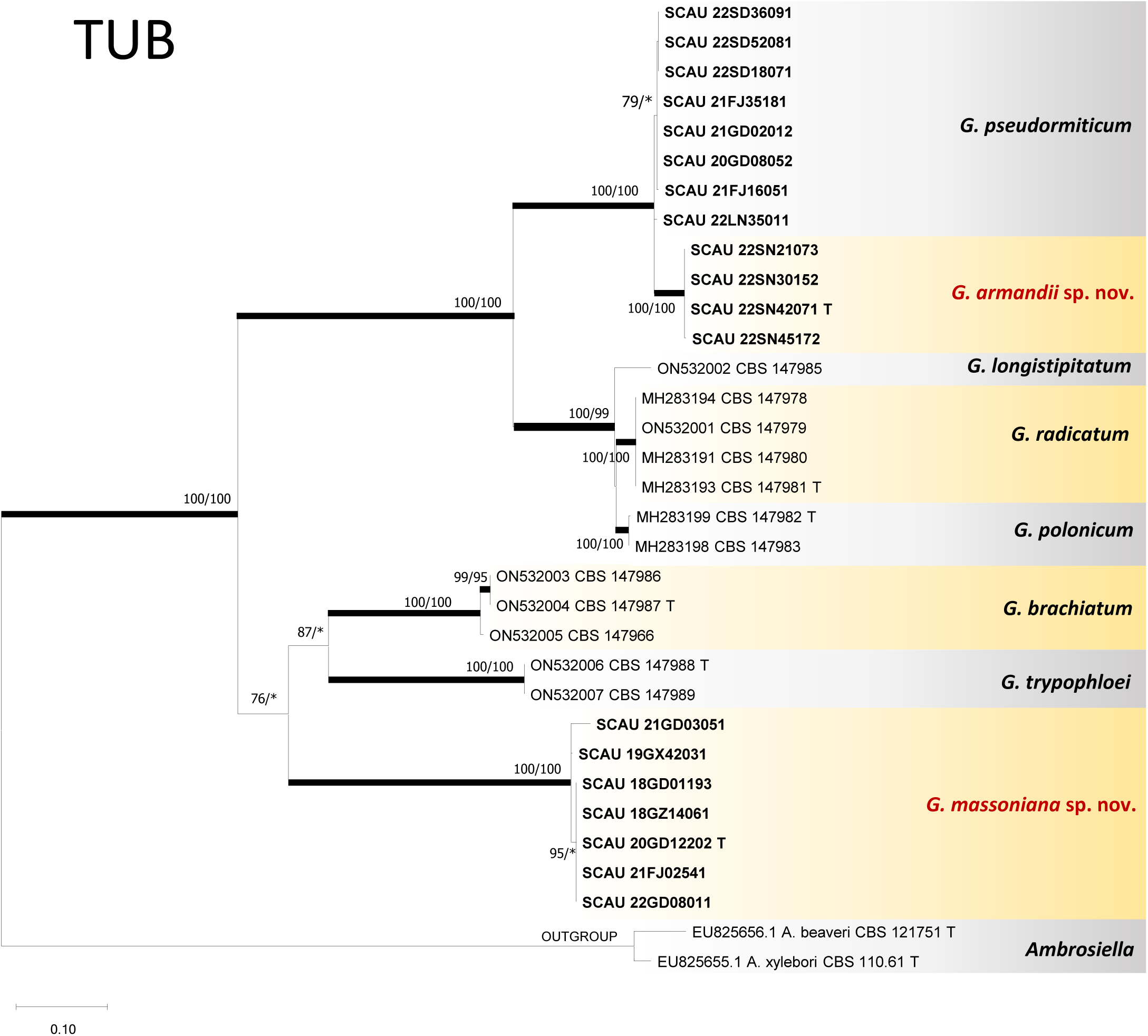
ML tree of *Graphium* generated from TUBB sequence data. Sequences generated from this study are printed in bold type. Bold branches indicate posterior probability values ≥ 0.95. Bootstrap values ≥75 % are recorded at nodes as ML/MP. * Bootstrap values < 75 %. T = ex-type isolates.

**Table 3.** Parameters used and statistical values related to all phylogenetic analyses in the present study.

|  |  | **ITS** | **LSU** | **TEF-1α** | **TUB** | **Combined** |
| --- | --- | --- | --- | --- | --- | --- |
| **Alignments** | **Taxa** | **72** | **46** | **58** | **33** | **37** |
|  | **Characters** | **576** | **529** | **375** | **687** | **1457** |
|  | **Constant** | **329** | **401** | **232** | **308** | **1016** |
| **MP** | **Informative** | **232** | **124** | **132** | **356** | **393** |
|  | **Uninformative** | **15** | **4** | **11** | **23** | **48** |
|  | **Tree number** | **48** | **3** | **2** | **3** | **1** |
|  | **Tree length** | **593** | **197** | **526** | **1082** | **902** |
|  | **CI** | **0.7352** | **0.8173** | **0.6844** | **0.8494** | **0.8137** |
|  | **RI** | **0.9512** | **0.9530** | **0.9335** | **0.9623** | **0.9463** |
|  | **RC** | **0.6994** | **0.7789** | **0.6389** | **0.8173** | **0.7701** |
|  | **HI** | **0.2648** | **0.1827** | **0.3156** | **0.1506** | **0.1862** |
| **Model tests** | **Sub. Models** | **GTR+I+G** | **GTR+I+G** | **GTR+I+G** | **GTR+I** | **TrN+I+G** |
| **ML** | **P-inv** | **0.6434** | **0.3701** | **0.5369** | **1.5732** | **0.3487** |
|  | **Gamma** | **0.2856** | **0.3483** | **1.9256** | **0.2242** | **0.4401** |

The ITS data matrix contained 17 known species of *Graphium* (Fig. 1). Four Shannxi isolates clustered as Taxon 1 under the branches of *G. fimbriisporum*, *G. pseudoumiticum,* and *G. laricis*. These isolates shared identical sequences with *G. pseudoumiticum*. Taxon 2, which consists of six isolates, was identified as a distinct lineage within the species complex, including *G. basitruncatum*, *G. carbonarium*, *G. euwallaceae,* and *G. kuroshium*. Eight isolates from this study in Taxon 3 were identical to *G. pesudomiticum* in the ITS gene region.

The LSU data matrix contained 12 known species but lacked *G. basitruncatum*, *G. fructicola*, *G. jumulu*, *G. putredinis,* and *G. scolytodis*. BI, ML, and MP analyses resulted in similar tree topologies (Fig. 2). Taxon 1 could be slightly different from *G. pseudoumiticum* (Taxon 3), while Taxon 2 formed a lineage from *G. carbonarium* and *G. jumulu* without high bootstrap support.

The aligned EF1A data matrix contained 15 known species, including exon 3 (sites 1-27), intron 3, exon 4 (sites 56-128), intron 5, and part of exon 5 (sites 325-427). Both BI and MP analyses agree with the results of ML analyses (Fig. 3). Taxa 1-3 and 15 known species were all well-defined. Taxon 1 was differentiated from *G. pseudoumiticum* (Taxon 3) and *G. laricis*. Meanwhile, Taxon 2 formed a separate lineage from G. basitruncatum, G. carbonarium, G. euwallaceae, and *G. kuroshium*.

The aligned TUBB data matrix contained exon 3 (sites 1 – 25), intron 3, exon 4 (sites 124 – 166), intron 4, exon 5 (sites 371 – 424), intron 5, and part of exon 6 (sites 592 – 687). The MP, BI, and ML resulted in similar tree topologies (Fig.4). All taxa and known species were well-defined. Although TUB sequences were not available, we identified this taxon as *G. pseudoumiticum* based on analyses of the other three genes.

### Taxonomy

*Graphium armandii* J. Li & M.L. Yin **sp. nov.** (Fig. 5)

**Figure 5.**
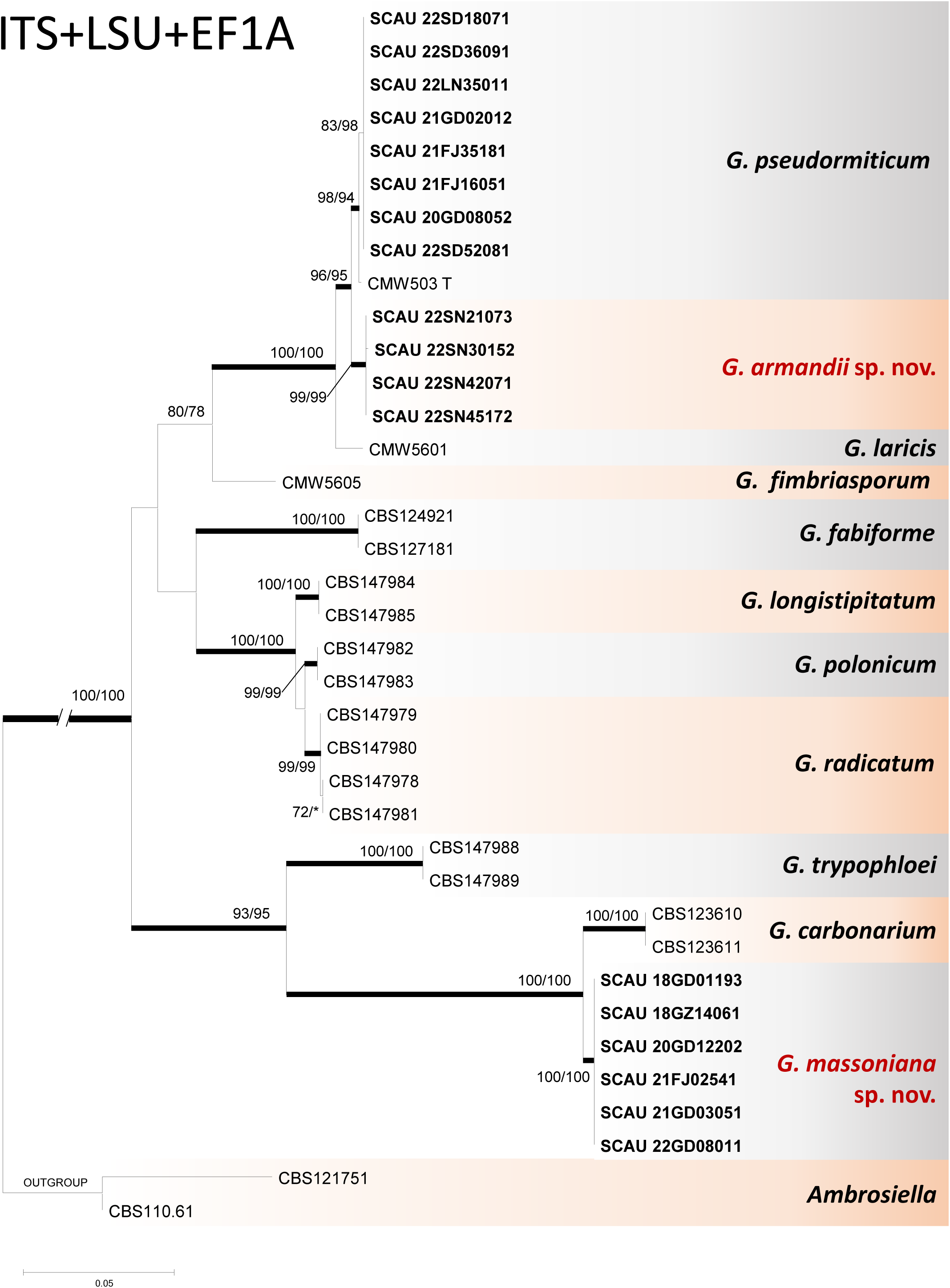
ML tree of *Graphium* generated from combined (ITS, LSU, EF1A) sequence data. Sequences generated from this study are printed in bold type. Bold branches indicate posterior probability values ≥ 0.95. Bootstrap values ≥75 % are recorded at nodes as ML/MP. * Bootstrap values < 75 %. T = ex-type isolates.

Mycobank: MB850358

Etymology. The name reflects the host tree, *Pinus armandii*, from which it was first found.

**Typus**. CHINA, Shannxi Province, Ankang, from the *Dendroctonus armandi* gallery in *Pinus armandii*, Aug. 2022, *M.J. Chen & M.L. Yin*, **holotype** HAMS 352758, ex-holotype culture SCAU 22SN42071 = CGMCC 3.27003.

Description. No sexual state was observed. Conidiophores macronemata, synnemata, abundant on MEA and sterilized twigs, individually or in clusters, expanding branches at the apex, (135–) 150 –161 (–166) μm long including conidiogenous apparatus, and (8–) 15 – 19 (–22) μm in width; Stipes brown to dark brown, wider at the top and narrower at the base, (80–) 95–110 (–133) in length, (13–) 15–18 (–20) μm in width; Rhizoids present, pigmented, septate, 18-30 in number, (80–) 93–115 (–120) in length; Conidiogenous cells annellated, (8–) 11 – 23 (–25) μm in length and (1–) 1.5 – 1.7 (–2) μm in width; Conidia aseptate, hyaline, cylindrical, smooth, slightly wider at the top and truncated at the base, (4–) 5 – 7 (–8.1) × (1.4–) 1.5 – 1.8 (–1.9) μm.

Culture Characteristics. The colonies in MEA were initially hyaline, later turning ivory-white to light yellowish, with rough and corrugated edges. The texture was dry and easy to lift. Hyphae were hyaline, appressed, and immersed. The optimal growth temperature for this organism is 25 °C, and its radial growth rate is 15 (± 0.5) mm/d. Growth is reduced at 10 °C and 30 °C and stops at 35 °C.

Additional specimens examined. CHINA, Shannxi Province, Ankang, from the gallery of *D. armandi* in *P. armandii*, Aug. 2022, *M.J. Chen & M.L. Yin*, SCAU 22SN21073, SCAU 22SN30152, and SCAU 22SN45172.

Insect vector: *D. armandi*

Host range: *P. armandii*

Distribution: China

Note: This species is phylogenetically close to *G. pseudormiticum*. However, they can be distinguished by the size of the conidia. The conidia of *G. armandii* are shorter than those of *G. pseudormiticum*.

*Graphium massoniana* J. Li & M.L. Yin **sp. nov.** (Fig. 6)

**Figure 6.**
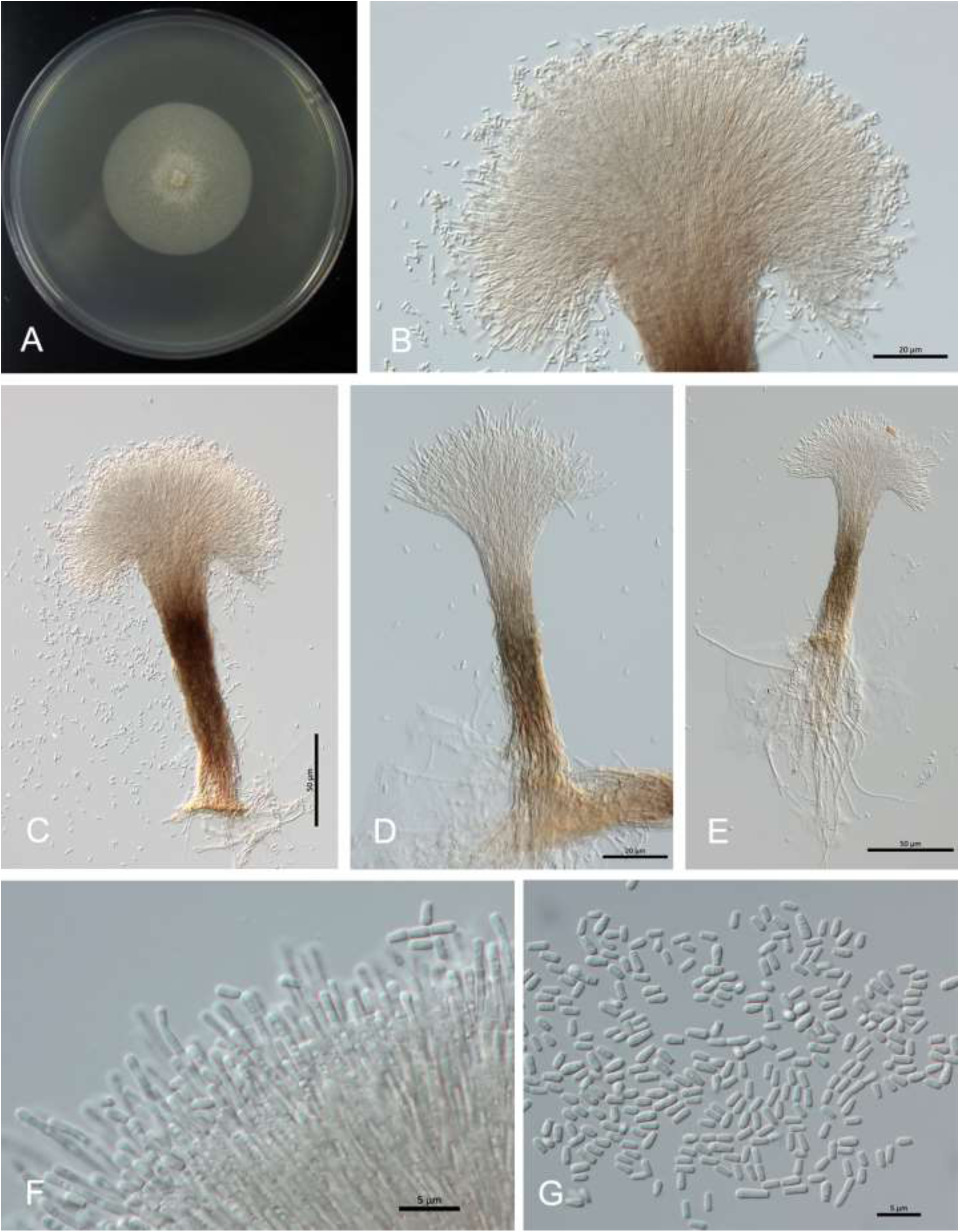
*Graphium armandii* sp. nov. (**A**) 14-day culture on MEA; (**B-E**) Synnematous asexual state in WA; (**F**) Conidiogenous apparatus; (**G**) Conidia

Mycobank: MB850359

Etymology. The epithet refers to the host tree, *Pinus massoniana*, from which it was first collected.

**Typus**. CHINA, Shannxi Province, Ankang, from the *Dendroctonus armandi* gallery in *Pinus armandii*, Aug. 2020, *X. Gao & M.L. Yin*, **holotype** HAMS 352757, ex-holotype culture SCAU 20GD12202 = CGMCC 3.27004.

Description. No sexual state was observed. Two types of Conidiophores: (i) stocky synnemata, macronemata, abundant in MEA and sterilized twigs, arising either individually or in clusters, expanding branches at the apex, pigmented, (90–) 105 –128 (–143) μm long including conidiogenous apparatus, (47–) 60 – 73 (–86) μm wide at the apex, (7–) 10 – 24 (–30) μm wide in the center, and 25 μm wide at the base; Stipes brown to light brown; Rhizoids-like hyphae present at the base; Conidiogenous cells 2-3 per branch point, annellidic conidiogenesis, (24–) 26 – 30 (–32) × 1.6 – 1.7 μm; Conidia aseptate, hyaline, cylindrical, smooth, slightly wider at the top and truncated at the base, 4.4 – 6.9 × 2.1 – 2.8 μm. (ii) slender synnemata, abundant on MEA and sterilized twigs, individually or occasionally in clusters, expanding branches at the apex, hyaline to olive brown, (180–) 215 – 273 (–400) μm long including conidiogenous apparatus, (16–) 20 – 26 (–33) μm at the apex, (6–) 7.5 – 9 (–11) μm wide in the center, and XX at the base; Stipes olive to olive brown, apex olive gray; Rhizoids-like hyphae absent; Conidiogenous cells annellated, 2-3 per branch point, (20–) 25 – 28 (–30) × 1.5 – 1.8 μm, Conidia are similar to that produced by former conidiophores.

Culture Characteristics. The colonies in MEA were initially hyaline and then milky white; the edges were intact and smooth, and the texture was dry and easy to lift. Hyphae are appressed and immersed in the agar. The optimal growth temperature is 25 °C and the radial growth rate is 31 (±0.5) mm/d. The growth rate decreases at 10 °C and 30 °C and stops at 35 °C.

Additional specimens examined. CHINA, Guangdong Province, Heyuan, from *Cyrtogenius luteus* gallery in *Pinus massoniana*, Oct. 2018, *X. Gao & M.L. Yin*, SCAU 18GD01193; Guizhou Province, Zunyi, from *Tomicus minor* gallery in *P. massoniana*, Aug. 2018, *X. Gao & M.L. Yin*, SCAU 18GZ14061; Guangxi Province, Liuzhou, from the *C. luteus* gallery *in P. massoniana*, Aug. 2019, *X. Gao & M.L. Yin*, SCAU 19GX42031; Fujian Province, Ningde, from the *Orthotomicus erosus* in *P. massoniana*, Aug. 2021, *X. Gao & M.L. Yin*, SCAU 21FJ02541, Guangdong Province, Zhaoqing, from the gallery of *O. erosus* in *P. massoniana*, Oct. 2021, *K. Liu & M.L. Yin*, SCAU 21GD03051; Guangdong Province, Qingyuan, from the gallery of *O. erosus* in *P. massoniana*, Oct. 2022, *K. Liu & M.L. Yin*, SCAU 22GD08011.

Insect vectors: *C. luteus*, *O. erosus*

Host range: *P. massoniana*

Distribution: China

Note: *G. massoniana* was phylogenetically placed in the clade, which includes *G. carbonarium*, *G. basitruncatum*, *G. euwallaceae*, and *G. jumulu* in this study. Morphological comparisons among these closed species were listed in Table **4**. Unlike other species with Scedosporium-like morphs, *G. massoniana* developed slender synnemata, making it unique regarding conidiophore formation. In addition, *G. massoniana* is the first species reported from pines associated with bark beetles in this clade.

**Table 4.**
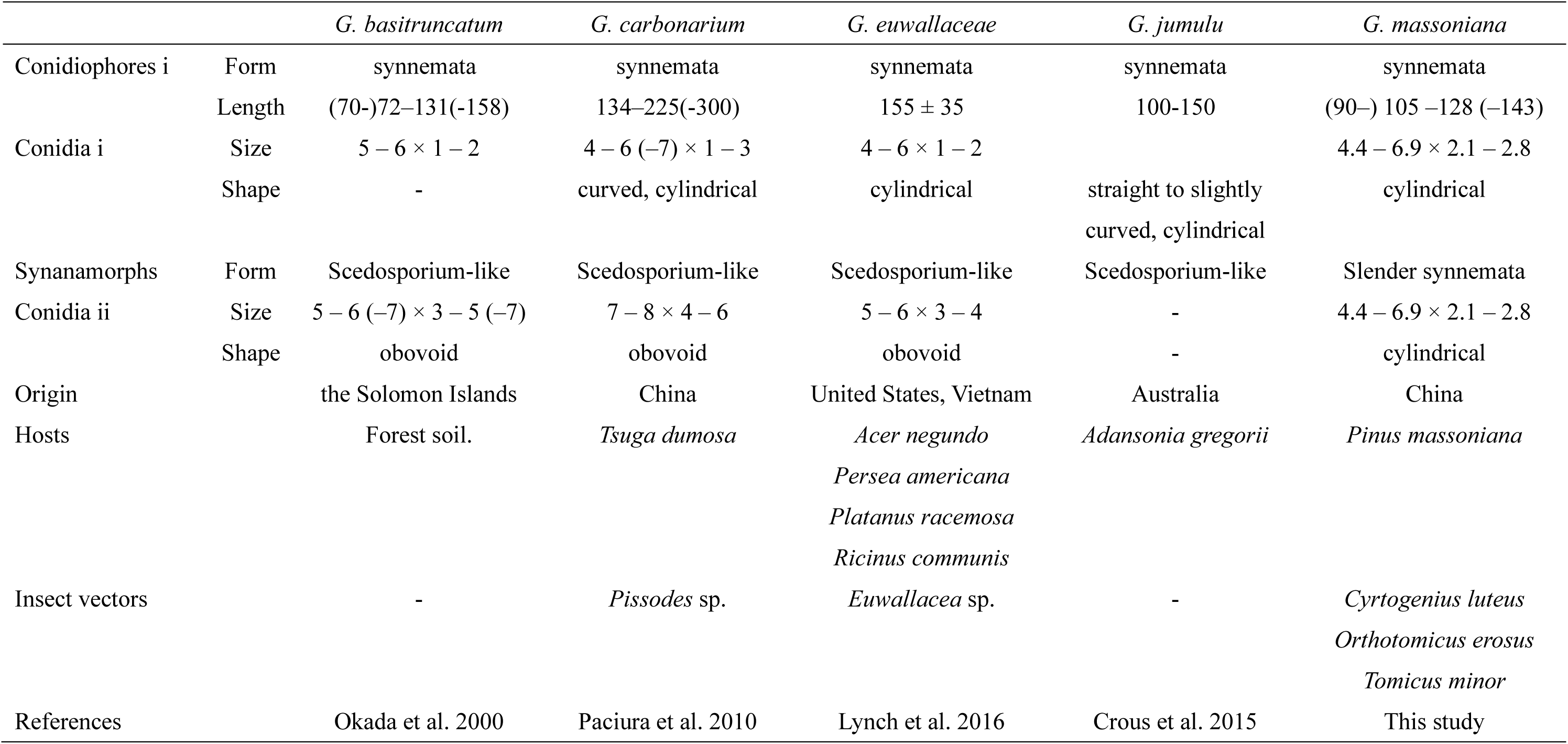
Comparisons of among *G. massoniana* sp. nov. and related species.

***Graphium pseudormiticum*** M. Mouton & M. J. Wingfield (Fig. 7)

**Figure 7.**
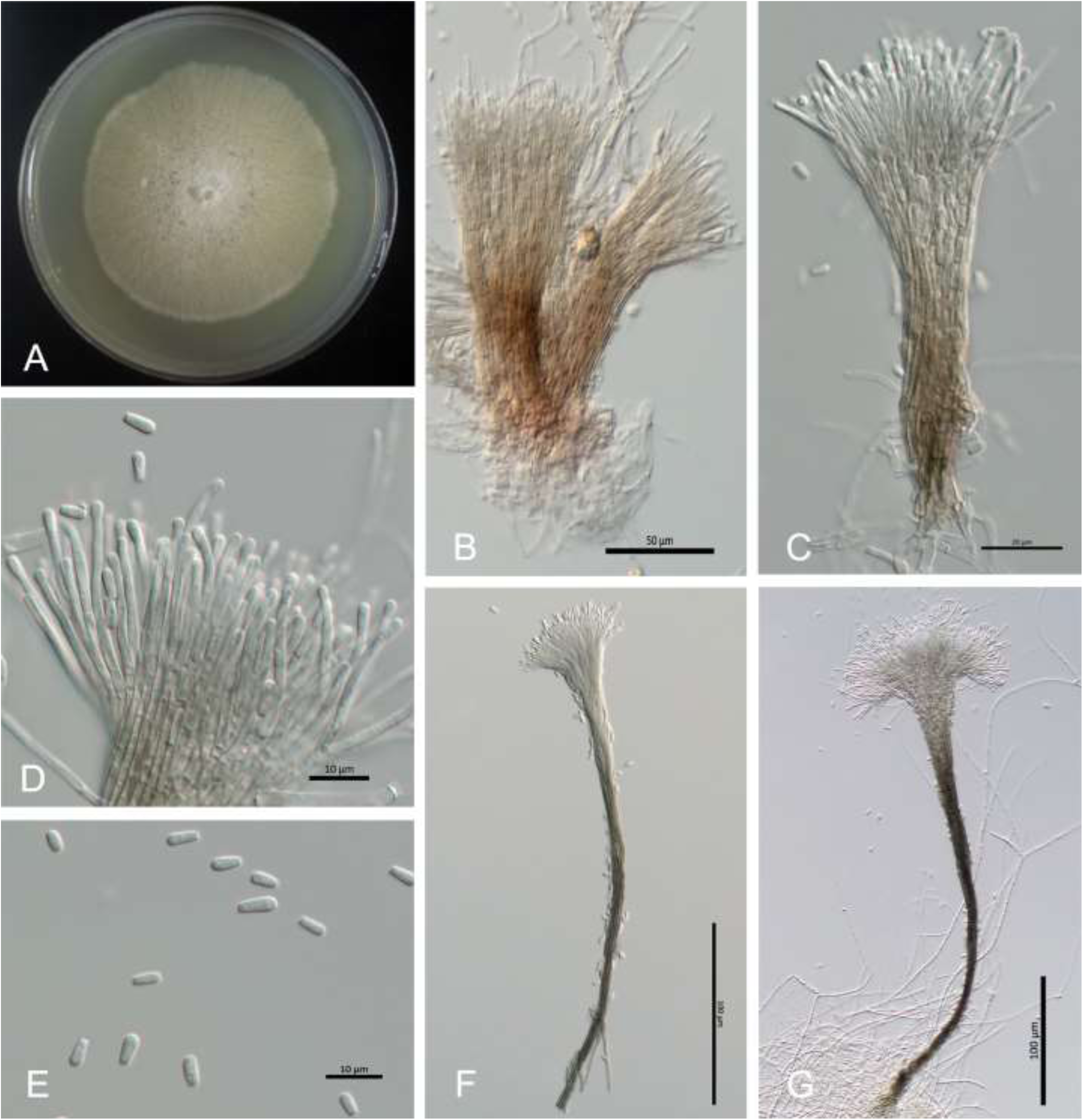
*Graphium massoniana* sp. nov. (**A**) fourteen-day-old culture on MEA; (**B-C**) Synnematous asexual state on WA; (**D**) conidiogenous apparatus; (**E**) Conidia; (**F, G**) Slender synnemata on WA.

**Figure 8.**
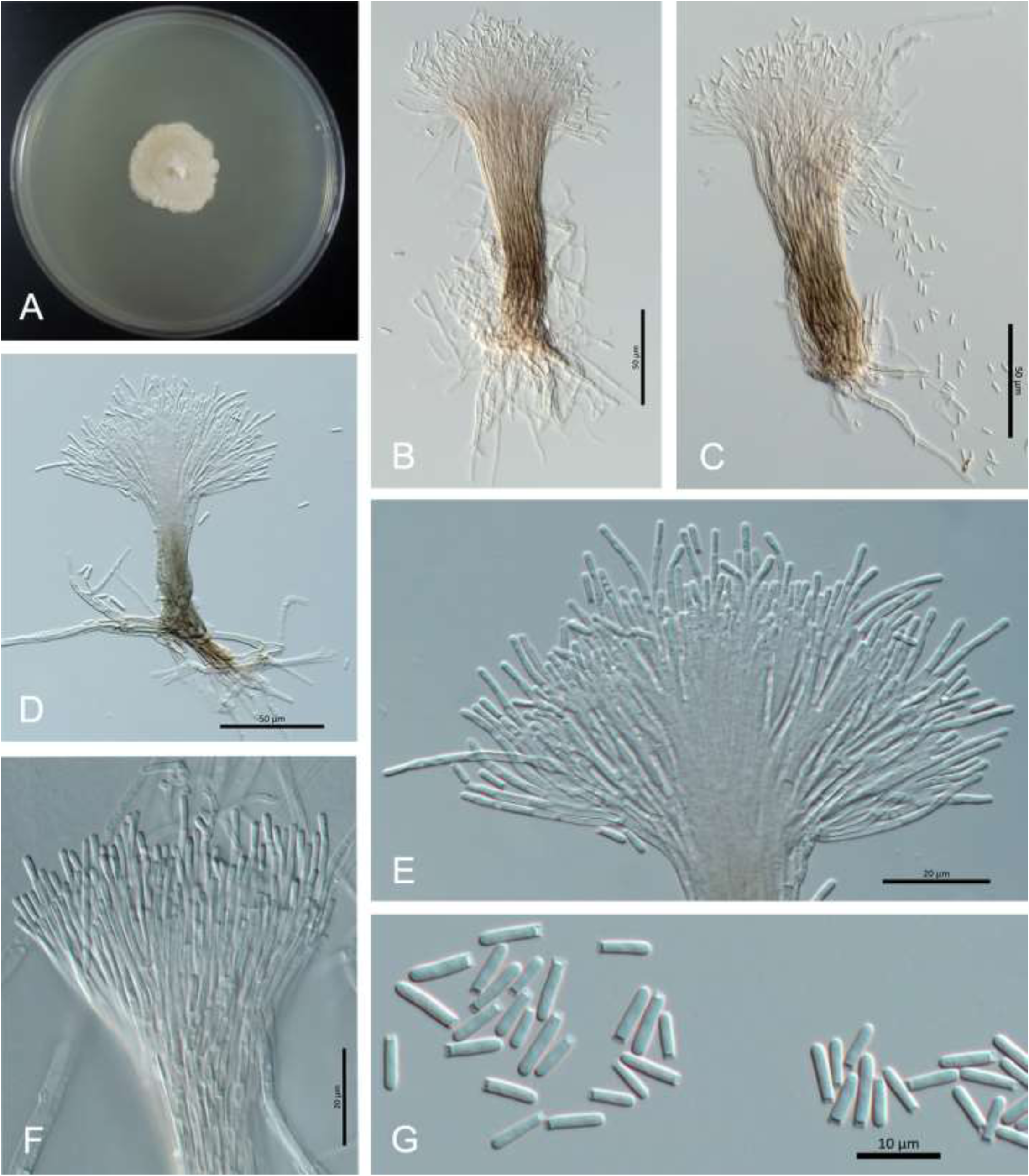
*Graphium pseudormiticum* (**A**) fourteen-day culture on MEA; (**B-D**) Synnematous asexual state in WA; (**E, F**) conidiogenous apparatus; (**G**) Conidia.

Description. Mouton et al. 1994 (Protologue)

Specimens examined in this study. CHINA, Guangdong Province, Shaoguan, from the *O. erosus* in *P. massoniana*; Oct. 2020, *X. Gao & M.L. Yin*, SCAU 20GD08052; Guangdong Province, Zhaoqing, from the gallery of *O. erosus* in *P. massoniana*, Oct. 2021, *K. Liu & M.L. Yin*, SCAU 21GD02012; Fujian Province, Ningde, from the *O. erosus* in *P. massoniana*, Aug. 2021, *X. Gao & ML. Yin*, SCAU 21FJ16051; Fujian Province, Sanming, from the gallery of *O. erosus* in *P. massoniana*, Aug. 2021, *X. Gao & M.L. Yin*, SCAU 21FJ35181; Shandong Province, Qingdao, from the *Cryphalus piceae* gallery in *Pinus thunbergii*, Aug. 2022, *C.W. Liu & M.L. Yin*, SCAU 22SD18071; Shandong Province, Yantai, from *Cr. piceae* gallery in *P. thunbergii*, Aug. 2022, *C.W. Liu & M.L. Yin*, SCAU 22SD36091; Shandong Province, Weihai, from *Cr. piceae* gallery in *P. thunbergii*, Aug. 2022, *C.W. Liu & M.L. Yin*, SCAU 22SD52081; Liaoning Province, Weihai, from the gallery of *Cr. piceae* in *P. koraiensis*, Aug. 2022, *Y.T. Ran & M.L. Yin*, SCAU 22LN35011.

Insect vectors: *Cryphalus piceae, Ips sexdentatus*, *Orthotomicus erosus*, *O. laricis, Tomicus minor*

Host range: *Pinus* spp., *P. massoniana*

Distribution: Austria, China, Germany, South Africa

Note: *G. pseudormiticum* was first reported in association with *O. erosus* in South Africa (Mouton et al. 1994). In this study, we found the same insect species in China. A new vector, *Cr. piceae*, and a new host, *P. massoniana,* have also been discovered. Based on our phylogenetic analyses, *G. pseudormiticum* is closely related to *G. armandii* and *G. laricis*.

## Discussion

*Graphium armandii* is currently only reported to be associated with *Dendroctonus armandi* in China. As an early-stage pest, *D. armandi* mainly damages healthy trees, and this native beetle caused significant damage in the southern part of Shaanxi and the northern part of Sichuan, specifically in the forests of *P. armandii* in the Qinling Mountains. (Wang et al. 2010). The symptoms of infestation include yellowing of the tree crown, which can lead to the tree’s death within 1-3 years.

On the contrary, *G. pesudomiticum* showed more diversified selections toward host trees and vector insects. In the present study, *G. pesudomiticum* was found to be associated with *O. erosus*, which infested *P. massoniana* in the southern region, and with *Cryphalus piceae*, which harms *P. tabuliformis* and *P. densiflora* in the northern region. This makes *G. pesudomiticum* the most widely distributed *Graphium* species in Chinese coniferous forests. Jacobs et al. (2003) believed that the host plants of *G. pseudormiticum* are limited to conifers. The feeding habits of insect vectors greatly influence the selection of host trees by associated fungi. Interestingly, this fungus was first reported to be associated with *O. erosus* in South Africa, the same insect vector in this study. *Orthotomicus erosus* is naturally distributed across Europe, North Africa, and Asia. It has been introduced in Chili, Fiji, South Africa, Swaziland, and the USA (Haack 2004). Pests commonly infest fallen or weakened trees, but can kill healthy trees when the population becomes large (Haack 2004). In China, this beetle is mainly found in *P. massoniana,* but can also infest *P. yunnanensis* and *P. kesiya* (Yin 1984).

Moreover, *Cryphalus piceae* can be found in central and southern Europe and parts of western Asia, where fir (*Abies*) and spruce (*Picea*) are abundant (Justesen et al. 2023). This beetle primarily infests branches of host trees, particularly those that are stressed, recently dead, or newly broken. Healthy trees are generally only at risk of attack when beetle populations have grown significantly (Cerchiarini et al. 1997). *Cryphalus piceae* was only known in China for infesting pines in Shandong (Chang et al. 2021).

As the second new species discovered in this study, *G. massoniana* can be carried by various types of bark beetles, *O. erosus* and *Cyrtogenius luteus*, and is widely distributed in Masson pine forests in southern China. However, other species in the *G. carbonarium* complex are found mostly in hosts of hardwood trees. For example, *G. basitruncatum* was reported as an opportunistic fungal pathogen in humans, which caused skin infection and fungemia symptoms in a patient with leukemia patient in Canada (Kumar et al. 2007). *G. carbonarium* was reported from Pissodes sp. on T. Dumosa in Yunnan (Paciura et al. 2010). *G. euwallaceae* (Lynch et al. 2016) and *G. kuroshium* (Na et al. 2018) were first described from *Euwallacea* sp. of infested avocado trees (*Persea americana*) in the USA. In western Australia, *G. jumulu* was reported from Adansonia gregorii (boab) (Crous et al. 2015). Research on *Graphium* fungi in hardwood trees in China is lacking. Therefore, it is reasonable to believe that more potential new species are still waiting to be reported from the country.

This study aims to fill the knowledge gaps in our understanding of *Graphium* associated with bark beetles in particular regions of Chinese pine forests. Further exploration of the role of *Graphium* species during beetle colonization in hosts and potential impacts on host trees is required.

## Acknowledgments

We thank all the forest staff who assisted us during the sampling process in Guangdong, Fujian, Guangxi, Guizhou, Shandong, Liaoning and Shannxi Provinces over the years, and we especially appreciate Professor J. Wang for his unwavering support and care for our laboratory-related work.

## Declarations

### Availability of data and material

The manuscript mentioned all the availability of data. Ex-type cultures of new species were deposited in the Culture Collection of South China Agricultural University (SCAU) and the China General Microbiological Culture Collection Center (CGMCC). The type herbariums were preserved in the Fungarium (HAMS), Institute of Microbiology, Chinese Academy of Sciences. DNA sequence data are available in Genebank (https://www.ncbi.nlm.nih.gov/nucleotide/), and taxonomic novelties are available in Mycobank (https://www. mycobank.org).

### Competing Interests

The authors declare no conflicts of interest.

### Funding

This work was funded by the National Natural Science Foundation of China (32070012) and the Guangdong Basic and Applied Basic Research Foundation (2020A1515010486, 2022A1515010901).

### Authors’ contributions

Conceptualization, Mingliang Yin; Data curation, Jie Li; Funding acquisition, Mingliang Yin; Formal analysis, Jie Li; Funding acquisition, Mingliang Yin; Investigation, Xiang Gao, Kun Liu, Minjie Chen, Yutong Ran and Congwang Liu; Methodology, Mingliang Yin; Project administration, Mingliang Yin; Resources, Xiang Gao, Kun Liu, Minjie Chen, Yutong Ran and Congwang Liu; Software, Jie Li; Supervision, Mingliang Yin; Validation, Mingliang Yin; Visualization, Mingliang Yin; Writing – original draft, Jie Li; Writing – review & editing, Tong Lin and Mingliang Yin.

## Notes

### Competing Interest Statement

The authors have declared no competing interest.

